# SRSA-VAE: Self-Attention-Based Feature Learning for Single-Cell Multimodal Clustering

**DOI:** 10.64898/2026.05.06.723212

**Authors:** Rangan Das, Ashmita Dey, Ujjwal Maulik, Sanghamitra Bandyopadhyay

## Abstract

Clustering plays a critical role in the analysis of single-cell omics data for identifying cellular heterogeneity and uncovering biological mechanisms. However, the high dimensionality, sparsity, and multimodal nature of single-cell datasets such as single-cell RNA sequencing (scRNA-seq) and Cellular Indexing of Transcriptomes and Epitopes by Sequencing (CITE-seq) pose significant challenges for effective feature learning and representation learning. Traditional dimensionality reduction methods often rely on linear transformations and fail to capture complex nonlinear relationships between gene and protein expression profiles. In this work, we propose SRSA-VAE, a scalable variational autoencoder framework that integrates a residual self-attention encoder for context-aware feature learning and multimodal representation learning. The proposed model dynamically contextualizes gene and protein representations through a self-attention mechanism, enabling the encoder to capture inter-cell relationships and emphasize biologically informative signals. A scalable residual connection further stabilizes training and preserves essential input information during latent representation learning. We evaluate SRSA-VAE on five large-scale publicly available single-cell datasets, including both scRNA-seq and CITE-seq data, and compare its performance with established deep generative models. Experimental results demonstrate that SRSA-VAE consistently outperforms existing methods in Adjusted Rand Index (ARI) across benchmark datasets, with particularly strong gains on complex immune cell populations. Ablation studies further confirm the importance of the self-attention mechanism and residual connection in enhancing model stability and clustering accuracy. The proposed model offers a generalizable, robust, and scalable solution for single-cell clustering tasks.

**Code Repository:** https://github.com/rangan2510/srsa-vae

## I. Introduction

Single-cell omics technologies have revolutionized the study of cellular heterogeneity by enabling the measurement of gene expression and protein abundance at single-cell resolution. In particular, single-cell RNA sequencing (scRNA-seq) and Cellular Indexing of Transcriptomes and Epitopes by Sequencing (CITE-seq) provide complementary views of cellular states by capturing both transcriptomic and proteomic information. These technologies have become essential tools for identifying novel cell types, understanding developmental processes, and investigating disease mechanisms. However, the high dimensionality, sparsity, and technical noise inherent in single-cell datasets present significant challenges for downstream analysis, particularly for clustering and accurate identification of cell populations.

Recent advances in deep generative models, such as scVI, LDVAE, and totalVI, have demonstrated strong potential for modeling single-cell data by learning low-dimensional latent representations that capture biological variability while accounting for technical noise. Despite their success, several challenges remain. First, most existing encoder architectures treat features independently and do not explicitly model complex dependencies among genes and proteins, potentially overlooking contextual structure critical for distinguishing subtle cellular states. Second, high-dimensional single-cell data often contain redundant or noisy features that can degrade clustering performance if not properly weighted during representation learning. Third, many models struggle to scale efficiently to large datasets while maintaining stable training dynamics. These limitations motivate the development of more flexible architectures capable of modeling feature interactions while maintaining computational scalability.

To address these challenges, we propose SRSA-VAE, a scalable variational autoencoder framework that integrates a residual self-attention encoder for multimodal representation learning. The self-attention mechanism contextualizes gene and protein signals across cells, allowing the model to emphasize biologically informative structure while suppressing noise, and residual connections stabilize training and preserve important feature information during latent representation learning. The main contributions of this work are as follows:

- Novel architecture: We propose SRSA-VAE, a variational autoencoder framework that integrates residual connections with a self-attention encoder to model context-dependent relationships in multimodal single-cell data.
- Context-aware feature learning: The proposed self-attention mechanism dynamically captures context across cells and modalities, improving representation learning and clustering performance.
- Scalable multimodal analysis: The framework efficiently integrates scRNA-seq and CITE-seq modalities while maintaining computational scalability.
- Comprehensive evaluation: Extensive experiments on multiple public datasets demonstrate that SRSA-VAE outperforms existing models such as scVI, LDVAE, and TotalVI in clustering accuracy and convergence efficiency.

The remainder of this paper is organized as follows. Section II reviews related work on deep generative models for single-cell analysis. Section III describes the datasets, preprocessing pipeline, and the proposed SRSA-VAE framework with its architectural components. Section IV presents the experimental results, ablation studies, and comparisons with existing methods. Finally, Section V concludes the paper and outlines potential future research directions.

## II. Related Works

To address the limitation of the traditional dimension reduction methods, several variational autoencoder (VAE)-based models have been proposed. scVI [1] introduced a deep generative model that performs probabilistic inference over single-cell RNA-seq data. LDVAE [2] further simplified the scVI architecture with a linear decoder to improve interpretability [3], [4]. For multimodal data (e.g., RNA and surface proteins), TotalVI [5] extended scVI to jointly model gene and protein expression [6].

Recent transformer-based frameworks such as scGPT [7] and contrastive learning approaches like SCCCL [8] have demonstrated strong generalization capabilities across large-scale single-cell datasets. On the other hand, for multimodal integration, MultiVI [9] extends the scVI framework to jointly model chromatin accessibility and gene expression, enabling unified embeddings. *BindSC* [10] applied manifold alignment to integrate cross-modal features such as RNA and ATAC data. However, these models are designed as foundation or pretraining frameworks that require substantial computational resources and large reference atlases. In contrast, SRSA-VAE is a lightweight generative model focused specifically on interpretable feature learning and clustering within individual datasets, making it methodologically distinct and complementary rather than directly comparable.

While recent models represent major advances in single-cell data integration, several challenges remain unresolved. Most existing frameworks either rely on modality-specific encoders or apply uniform transformations to all features, overlooking contextual dependencies among genes and proteins. Moreover, transformer-based models such as scGPT require extensive pre-training and computational resources, making them less accessible for medium-scale biological studies. Therefore, there is a continued need for lightweight, interpretable, and scalable models capable of capturing nonlinear, cross-modal relationships with minimal training cost. SRSA-VAE is developed precisely to fill this gap.

## III. Materials and Methods

This section discusses the method for dataset preprocessing, followed by the embedding and clustering process. The overall encoder architecture is summarized in Figure 1.

**Fig. 1.**
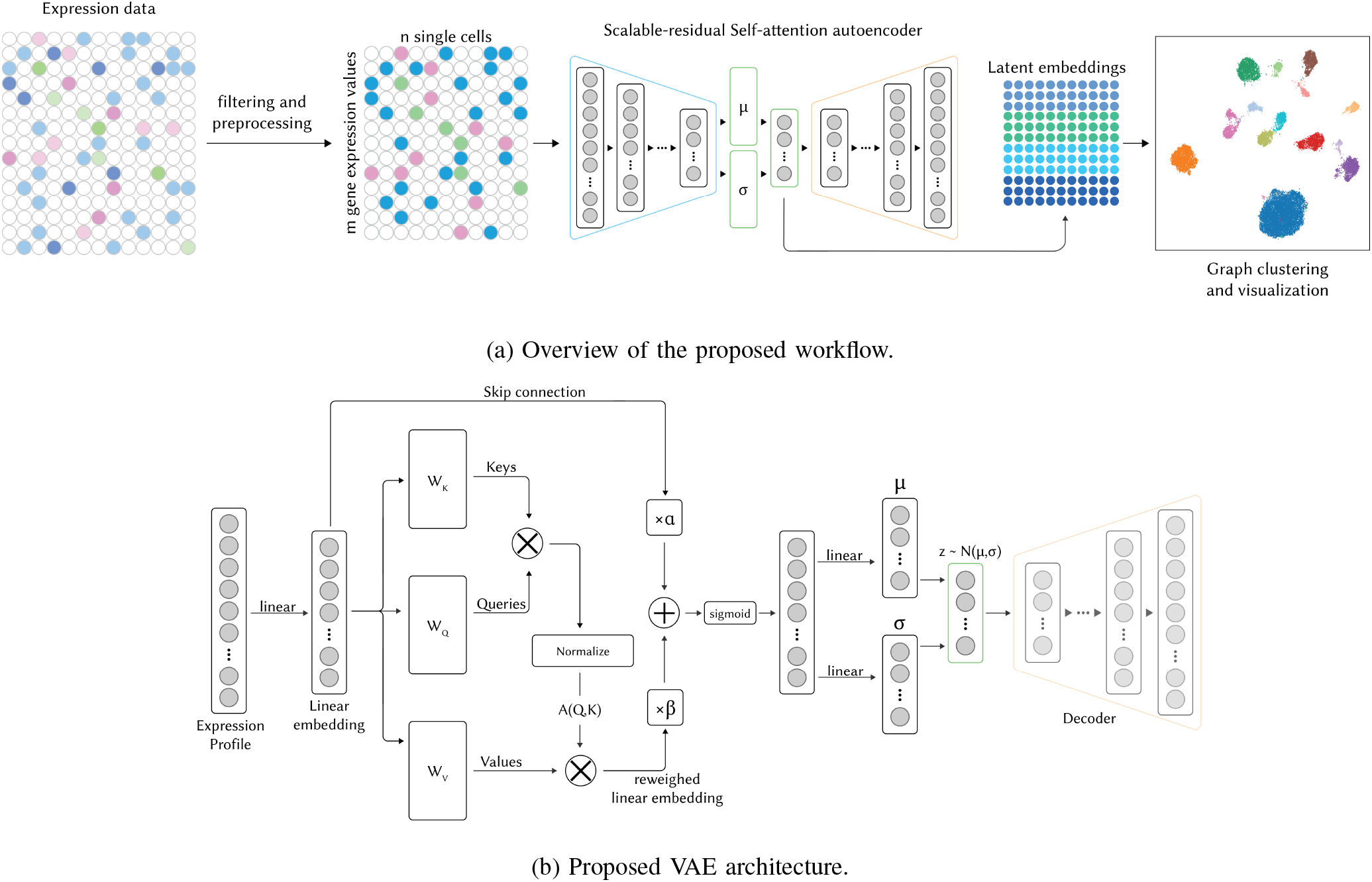
(a) Overview of the proposed workflow. (b) Internal structure of the SRSA-VAE encoder. Given the expression profile *X*, the encoder first generates a linear embedding corresponding to meta-genes. The embedding is then reweighted using a self-attention mechanism, keeping the dimensionality unchanged. The original linear embedding, scaled by a learnable parameter *α*, and the attention-reweighted embedding, scaled by *β*, are combined via element-wise summation (+) and passed through a sigmoid activation within the residual gate. Symbols (×) denote scalar multiplication by the learnable parameters *α* and *β*.

### A. Real Datasets

In order to evaluate the proposed self-attention-based feature learning mechanism for cell-type clustering, we use five real publicly available single-cell datasets that include both scRNA-seq and CITE-seq data. In Table I, the primary information of the real datasets is reported. Litvinukova et al. [11] highlighted the transcriptome of 486,134 cells and nuclei from six anatomical cardiac regions. Gayoso et al. [5] demonstrated RNA and surface protein abundance in single cells with CITE-seq from the spleen and lymph nodes of two wild-type mice. Hao et al. [12] applied their procedure to a CITE-seq dataset of 211,000 human peripheral blood mononuclear cells (PBMCs) with panels extending to 228 antibodies to construct a multi-modal reference atlas of the circulating immune system; after quality-control filtering, 161,764 cells were retained for our experiments. The PBMC 12K dataset [13] consists of scRNA-seq data from two batches of peripheral blood mononuclear cells from a healthy donor (4K PBMCs and 8K PBMCs). PBMC 68K [14] consists of ∼ 68,000 PBMCs collected from a healthy donor. Single-cell expression profiles of 11 purified subpopulations of PBMCs are used as references for cell type annotation. This dataset served as a gold standard for performance assessment of the clustering techniques.

**TABLE 1.**
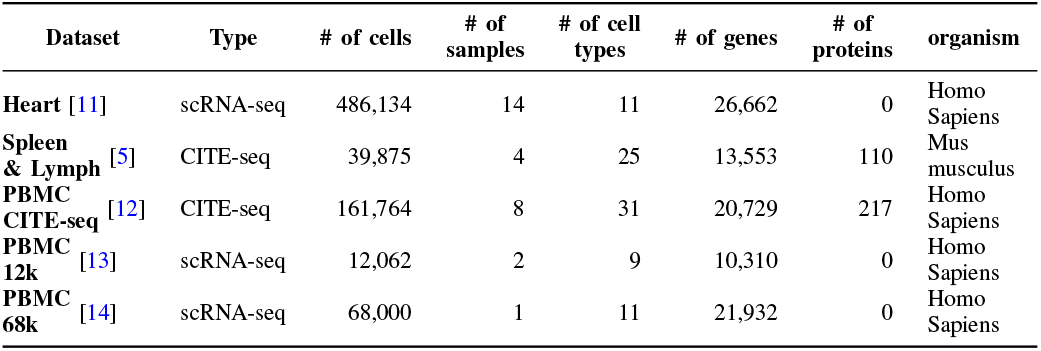
Specifications of the real datasets used in the paper for evaluation of the methods.

| Dataset | Type | # of cells | # of samples | # of cell types | # of genes | # of proteins | organism |
| --- | --- | --- | --- | --- | --- | --- | --- |
| Heart [11] | scRNA-seq | 486,134 | 14 | 11 | 26,662 | 0 | Homo Sapiens |
| Spleen & Lymph [5] | CITE-seq | 39,875 | 4 | 25 | 13,553 | 110 | Mus musculus |
| PBMC CITE-seq [12] | CITE-seq | 161,764 | 8 | 31 | 20,729 | 217 | Homo Sapiens |
| PBMC 12k [13] | scRNA-seq | 12,062 | 2 | 9 | 10,310 | 0 | Homo Sapiens |
| PBMC 68k [14] | scRNA-seq | 68,000 | 1 | 11 | 21,932 | 0 | Homo Sapiens |

### B. Data preprocessing

To preprocess the datasets, the proposed method follows the workflow in [15] with some additional changes for the proposed method. We filter out genes that are expressed in fewer than 3 cells. We also remove the cells that have fewer than 200 genes expressed. This reduces the sparsity of the expression matrix. Furthermore, for quality control, we remove the cells with *>* 5% of mitochondrial genes expressed. Next, we log-normalize the data and select 1, 200 highly variable genes using the pipeline proposed in Seurat v3 [15]. The ranking of the highly variable genes is performed by standardizing the data using z-score normalization per feature, followed by a regularized standard deviation. Following the regularization, the genes are ranked according to their normalized variance.

### C. Meta-Feature Generation with SRSA-VAE Encoder

We propose a feature learning framework using a variational autoencoder (VAE) with a scalable residual self-attention encoder (SRSAE). In classical VAEs, each input feature contributes equally to the posterior inference of latent variables, which assumes independence and ignores contextual structure in the data. This uniform treatment can obscure critical co-expression patterns that drive cellular heterogeneity. SRSA-VAE modifies this assumption by introducing a *context-adaptive posterior inference*, where each cell representation is dynamically informed by its relationships to other cells in the batch.

Concretely, the linear layer first maps each cell’s expression vector to **h**^(1)^ ∈ ℝ^*d*^ (see Section III-E1). Stacking all cells in a mini-batch gives 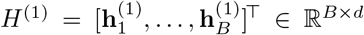. The encoder then computes a cell-context matrix:

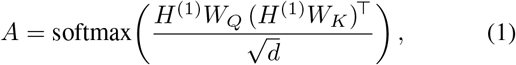

where *A* ∈ ℝ^*B*×*B*^ captures cross-sample affinities within the batch. The attention-refined representation is

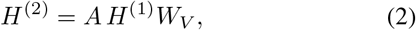

so that each cell’s embedding is contextually informed by other cells in the same mini-batch. Because the attention weights are computed dynamically from the batch content, the model naturally adapts to varying batch sizes and compositions; at inference time the full dataset (or any desired subset) is passed through the trained encoder in a single forward pass to obtain the final latent representations.

This formulation enables the model to capture nonlinear conditional dependencies across cells and modalities, effectively redefining the variational posterior *q*(**z**|*H*^(1)^) as a weighted function of biologically meaningful context rather than independent dimensions. For scRNA-seq-only datasets (those with no protein measurements), the same architecture is used with RNA features alone.

In addition, the *scalable residual connection* introduces a smooth interpolation between the initial linear embedding and its attention-refined transformation, represented as:

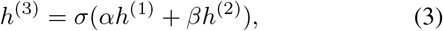

where *α* and *β* are trainable scaling factors. This parameterization yields a bounded fusion of the linear and attention-refined embeddings, ensuring model stability and preventing attention overfitting, a common issue in deep self-attention architectures.

To preserve input fidelity and stabilize training, a skip connection combines the original and attention-enhanced embeddings. Two trainable scalars *α* and *β* adjust the residual and attention contributions before final activation. SRSA-VAE consistently selects 6–10 latent features depending on dataset complexity and outperforms PCA in retaining cluster-relevant features, as shown in Figure 2.

**Fig. 2.**
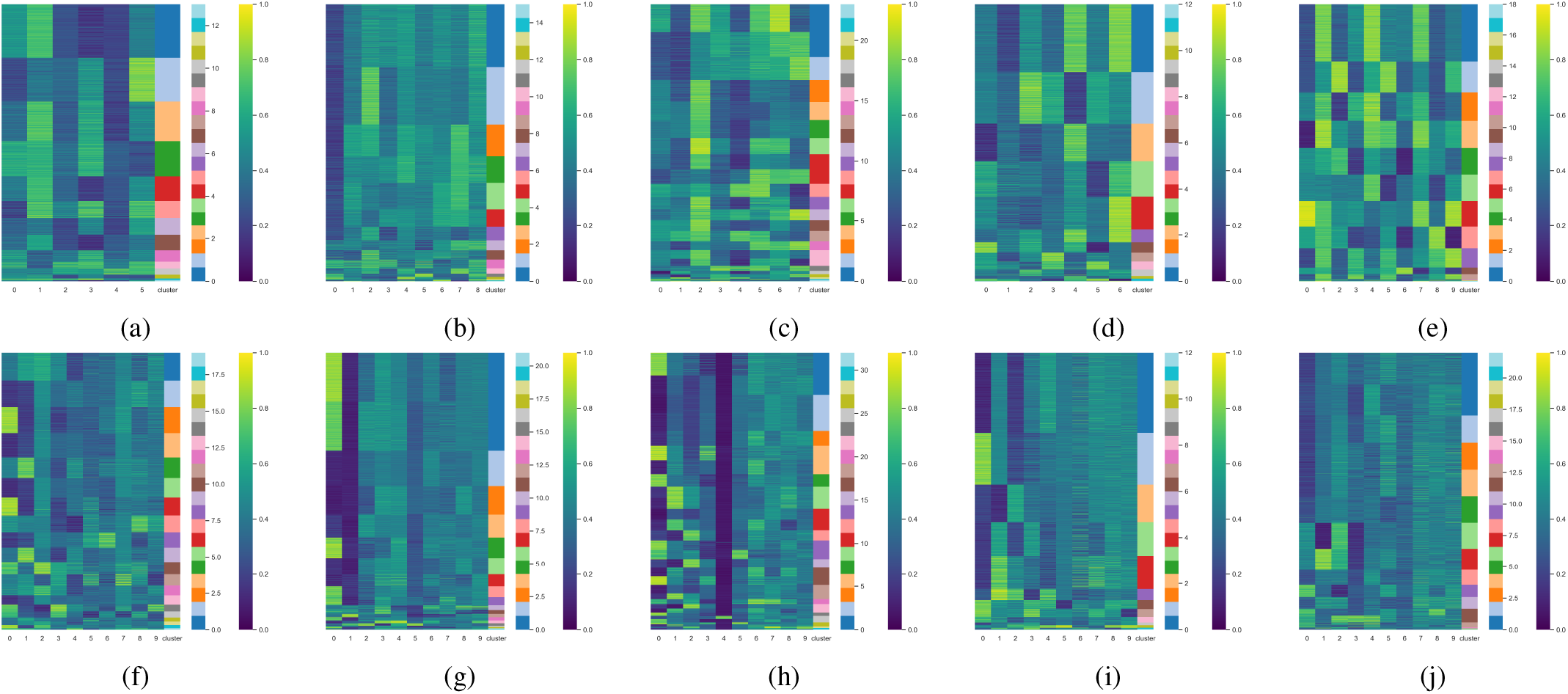
Cluster map of the latent variables and how they contribute to the clustering of the datasets Heart (a,f), Spleen and Lymph (b,g), PBMC CITE-seq (c,h), PBMC 12k (d,i) and PBMC 68k (e,j). The embedding of the proposed method is shown in subfigures (a)–(e), whereas the PCA embedding is shown in subfigures (f)–(j).

This design advances the theoretical basis of variational inference in single-cell analysis by embedding context-aware, feature-level adaptivity into the probabilistic learning process. SRSA-VAE thereby bridges the gap between statistical feature learning and deep generative modeling, offering a more interpretable and mathematically grounded solution for multimodal single-cell clustering.

### D. Variational Inference

This section discusses the inference of the latent embeddings. Given a gene expression profile *X* for a single cell, the model generates a vector *z* where *X* is a high-dimensional space and *z* is a low-dimensional embedding of *X*. Given *z* is a single latent encoding of a set of gene expression values for a single cell, a variational auto-encoder (VAE) tries to model the distribution *P* (𝒳) of the expression values in a high-dimensional space 𝒳. In the VAE model, the encoder network generates the samples *z* and the decoder network maps them to their original dimensions. The idea here is to generate a *z* that increases the probability of recovering the original gene expression profile. By this process, we can generate a low dimensional encoding that has the relevant biological signals.

To generate this *z*, we have to compute the posterior *P* (*z* |𝒳), which for a continuous latent variable model can be expressed as:

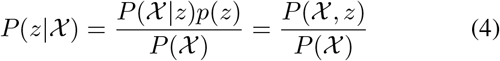

where *P* (*z*) and *P* (*z* |𝒳) denote continuous probability density functions of the latent variable and observed data, respectively. *P* (𝒳) represents the marginal likelihood obtained by integrating over all possible *z* values.

However, computing *P* (𝒳) is challenging and in many cases, it is intractable. Hence, the VAE tries to approximate the posterior using a variational probability *Q*(*z* | 𝒳) and Kullback–Leibler (KL) divergence to measure the dissimilarity between *Q* and *P*. This is given as:

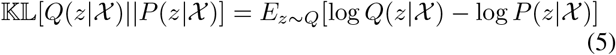

Using Bayesian inference, this can be rearranged as:

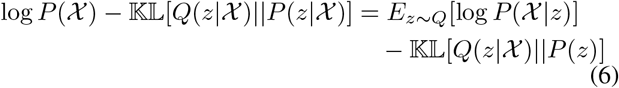

where log *P* (𝒳) is constant with respect to the variational parameters. *E*_*z*∼*Q*_ is the expectation over *z* sampled from *Q*. Therefore, it can be said that the problem of minimizing the KL divergence is the same as maximizing the right-hand side of (6). This can be performed using two neural networks with the ELBO as the training objective. The encoder approximates *Q*(*z* | 𝒳) and the decoder models *P* (𝒳 | *z*), or more simply, the encoder generates *z* from and the decoder tries to reconstruct it. By varying the structure of these neural networks, we can obtain different latent representations.

A Gaussian distribution is used to infer the latent variables with a normal prior 𝒩 (0, *I*). The encoder network estimates the posterior parameters *µ* and *σ* through dedicated linear projection layers, as detailed in Section III-E4.

### E. Scalable–Residual Self-Attention Encoder for Expression Imputation

To infer the posterior parameters of the latent variables, we introduce a novel encoder architecture that imputes biological signals in a reduced-dimensional space. Our implementation extends the scVI-tools framework; the architecture is illustrated in Figure 1b. This encoder achieves comparable or superior clustering performance with fewer latent dimensions and substantially reduces the number of training epochs required for convergence relative to the default scVI model. The following subsections detail each layer of the encoder network.

1. *Linear Transform:* We first apply dropout to the input expression vector **x** ∈ ℝ^*p*^ at rate 0.1, followed by a fully connected layer, batch normalization, and a ReLU activation. Denoting the input (after dropout) as **x**^(0)^, the layer output **h**^(1)^ ∈ ℝ^*d*^ (with *d* = 128) is

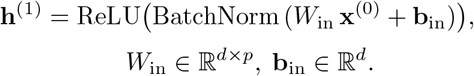
2. *Self-Attention:* The vector **h**^(1)^ is then processed by a single-head self-attention layer that re-weights its components. We compute

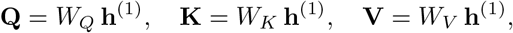

with *W*_*Q*_, *W*_*K*_, *W*_*V*_ ∈ ℝ^*d*×*d*^. The attention matrix *A* ∈ ℝ^*B*×*B*^ for a batch of size *B* is

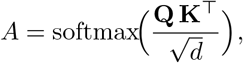

and the attended output **h**^(2)^ ∈ ℝ^*d*^ is

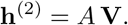 Note that the attention operates across cells within a batch rather than across features within a single cell. This cross-sample formulation allows each cell’s representation to be contextually informed by other cells in the batch, enabling the encoder to capture shared expression patterns that distinguish closely related cell types.
3. *Scalable Residual Connection:* To preserve the original meta-gene signals, we fuse **h**^(1)^ and **h**^(2)^ via a gated residual connection. Two trainable scalars *α, β* ∈ ℝ (initialized to 0.1 and 0.9) weight the contributions:

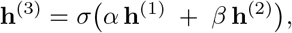

where *σ*(·) denotes the sigmoid activation and **h**^(3)^ ∈ ℝ^*d*^.
4. *Latent Sample Generation:* Finally, **h**^(3)^ is projected to the Gaussian variational parameters. Two linear layers predict

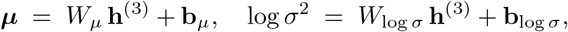

with *W*_*µ*_, *W*_log *σ*_ ∈ ℝ^*m*×*d*^ and **b**_*µ*_, **b**_log *σ*_ ∈ ℝ^*m*^. A latent sample **z** ∈ ℝ^*m*^ is then drawn via the reparameterization trick:

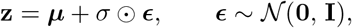

where 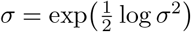 and ⊙ denotes element-wise multiplication.
5. *Decoder:* Our implementation reuses the default scVI decoder without modification. Given the latent sample **z** ∈ ℝ^*m*^ and a one-hot batch indicator, the decoder consists of a single fully connected hidden layer (128 units, batch normalization, ReLU activation), followed by output layers that parameterize a negative-binomial likelihood for gene expression counts and, when protein measurements are available, a negative-binomial mixture for protein counts. Full details of the generative model are described in [1], [5]; the encoder described above is the sole architectural modification introduced by SRSA-VAE.

### F. Training and Hyperparameter Optimization

The SRSA-VAE model is trained to optimize clustering performance on the learned latent space. Key hyperparameters include the number of training epochs and the dimension of the latent space **z**. We perform a grid search over the latent dimension to maximize two external clustering metrics: Adjusted Rand Index (ARI) [16] and Normalized Mutual Information (NMI) [17]. The effect of latent dimension on clustering accuracy is shown in Figure 5, while model convergence across epochs is illustrated in Figure 6. The training objective follows the standard variational autoencoder loss function:

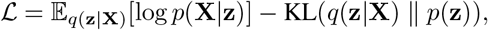

where the first term is the reconstruction loss and the second is the Kullback–Leibler (KL) divergence. We use the Adam optimizer with a learning rate of 0.001.

### G. Clustering

Let **X** ∈ ℝ^*n*×*d*^ denote the high-dimensional input data matrix, where *n* is the number of cells and *d* is the number of genes or features. After passing through the SRSA-VAE encoder, the model generates a low-dimensional latent representation **Z** ∈ ℝ^*n*×*k*^, where *k* ≪ *d*.

To identify distinct cell populations, we perform clustering on the latent space **Z**. Specifically, we construct a *k*-nearest neighbor (k-NN) graph 𝒢 = (𝒱, ℰ), where each node *v*_*i*_ ∈ 𝒱 corresponds to a cell embedding **z**_*i*_ ∈ ℝ^*k*^. The edges are formed based on Euclidean distances:

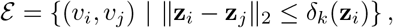

where *δ*_*k*_(**z**_*i*_) denotes the distance to the *k*-th nearest neighbor of **z**_*i*_.

We then compute a weighted adjacency matrix **W** ∈ ℝ^*n*×*n*^

where:

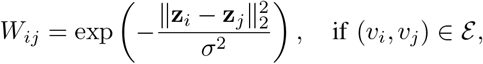

and *W*_*ij*_ = 0 otherwise. Here, *σ* is a scaling hyperparameter that controls local smoothness.

On the resulting graph 𝒢, we apply the Leiden community detection algorithm [18] to partition the nodes into *C* non-overlapping clusters {𝒞_1_, 𝒞_2_, …, 𝒞_*C*_}:

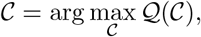

where 𝒬 denotes the modularity function used by the Leiden method to measure the quality of community assignments.

This graph-based clustering approach allows us to capture both local structure and global relationships in the latent space. The number of clusters *C* is determined by the resolution parameter of the Leiden algorithm. For a fair comparison, we applied the same fixed Leiden resolution of 0.5 to the embeddings produced by *all* methods (SRSA-VAE, scVI, LDVAE, and Scanpy/Seurat) across all five datasets. Under this uniform setting, the number of clusters identified by the Leiden algorithm varied across methods due to differences in latent-space geometry; for the SRSA-VAE embeddings specifically, the algorithm identified 14, 18, 24, 12, and 19 clusters for the Heart, Spleen & Lymph, PBMC CITE-seq, PBMC 12k, and PBMC 68k datasets, respectively.

1. *Evaluation criteria:* For the evaluation of the proposed method, we use both quantitative and qualitative analysis. For quantitative analysis, we focus on the clustering performance using the proposed embedding method. For qualitative analysis, we compare the UMAP embedding of the state-of-the-art methods as well as the proposed method. We use both intrinsic and extrinsic methods for the quantitative analysis of clustering performance. Given the ground truth is available for the dataset used in this paper, we measure the accuracy of the clustering through three extrinsic metrics - Adjusted Rand Index (ARI), Normalized Mutual Information (NMI) and V-measure [19] as shown in (7), (8), and (9), respectively.

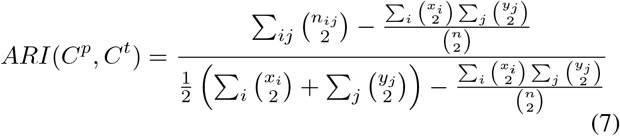

where *C*^*p*^ are the predicted clusters and *C*^*t*^ are the true clusters. *n*_*ij*_ are the total number of nodes that are present in 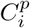 and 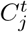. *x*_*i*_ is the total of all *n*_*ij*_ corresponding to any of *C*^*t*^ and all of *C*^*p*^. *y*_*j*_ is the summation of all *n*_*ij*_ corresponding to any *C*^*p*^ and all of *C*^*t*^.

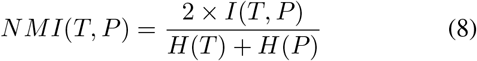

where T determines the true class labels, P is the predicted class labels, *H*(·) is the Entropy function and *I*(*X, Y*) is the mutual information between *X* and *Y*. V-measure is simply the harmonic mean of homogeneity and completeness and is given as:

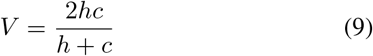

where *h* is the homogeneity score and *c* is the completeness score as described in [19]. For intrinsic clustering metrics, we use the Silhouette score and Calinski-Harabasz score. Silhouette score for a single point *i* is given as

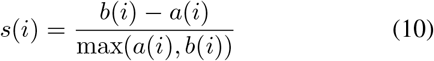

where *a*(*i*) is the mean distance between *i* and all other points in its own cluster and *b*(*i*) is the mean distance between *i* and all points in the nearest neighbouring cluster. Calinski-Harabasz score, which is the ratio between sum of inter-cluster dispersion and the sum of intra-cluster dispersion for all clusters, is given as

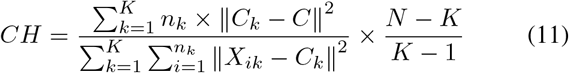

where *N* is the total number of samples, *K* is the total number of clusters, *n*_*k*_ is the number of observations in cluster *k, C*_*k*_ is the centroid of cluster *k, C* is the centroid of the dataset (barycenter) and *X*_*ik*_ is the *i*^*th*^ observation of cluster *k*.

## IV. Results

The following section discusses the effect of the proposed embedding method on the downstream clustering performance.

### A. Effect of Proposed Approach on clustering

In this section, we describe the results of the proposed embedding method by comparing the clustering performance with other methods. Here, we compare the performance by comparing the clustering accuracy and quality metrics, and also by visualizing the clusters. For CITE-seq datasets, we also made a further comparison of the proposed model with a model designed solely for such multimodal datasets.

#### Visualization of latent embeddings

Visualization of the latent embedding allows us to perform a qualitative analysis of the clustering performance. We tested SRSA-VAE on five large-scale benchmarking datasets and compared them with scVI and LDVAE. LDVAE is similar to scVI, but with a linear decoder. We also used Scanpy to perform principal component analysis. Using Scanpy, we try to reproduce the results of Seurat [15]. Since we use the pre-processing as documented in Seurat v3, the clustering outputs of Scanpy resemble that of Seurat, which is considered a standard method. In each case, we visualized the low-dimensional embedding using UMAP. The results of the visualization are shown in Figure 4. SRSA-VAE performs notably better across all the datasets when it comes to visualizing the clusters in a well-separated manner.

**Fig. 3.**
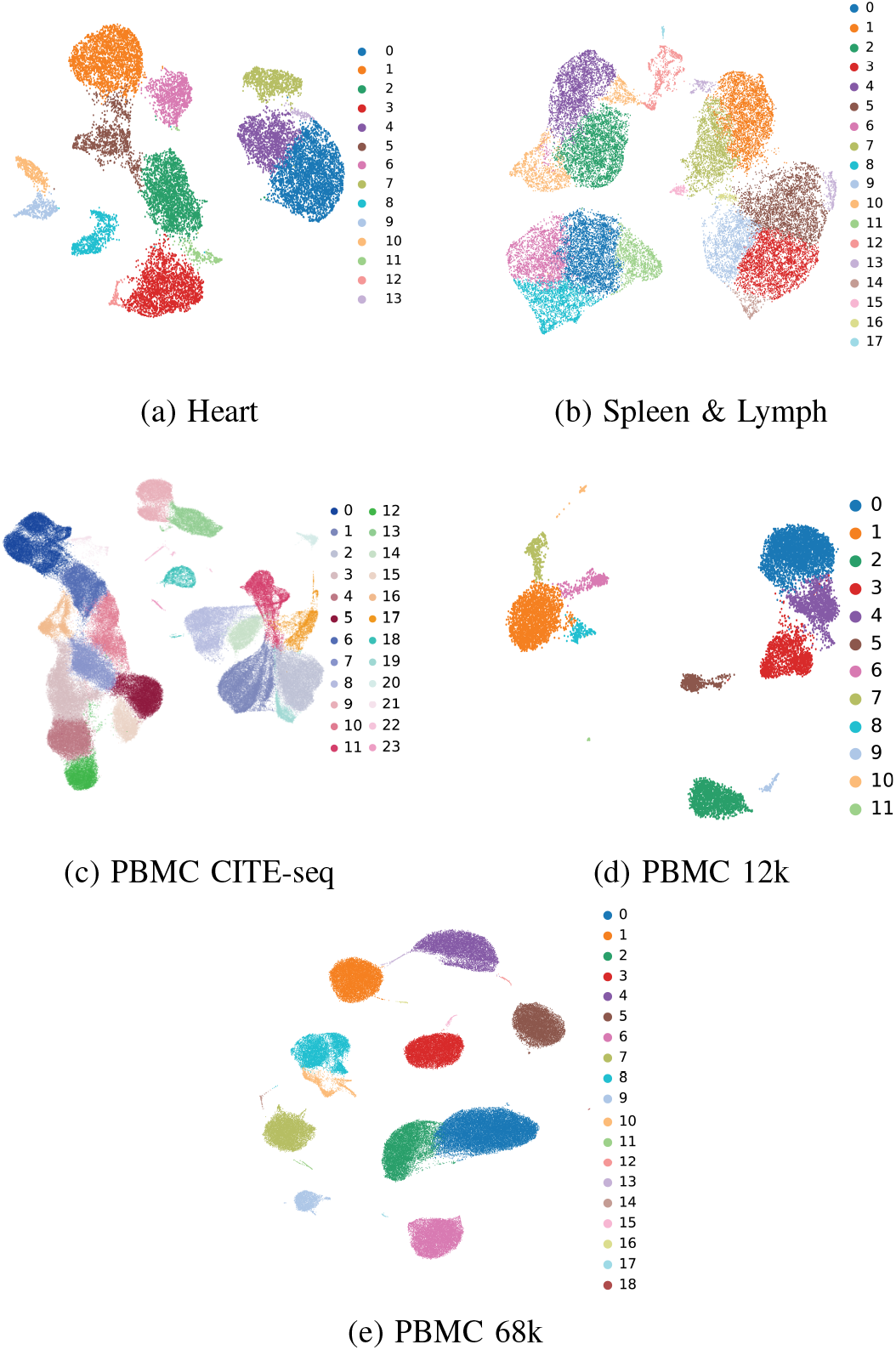
UMAP representation of clustering outputs on five datasets using the proposed method.

**Fig. 4.**
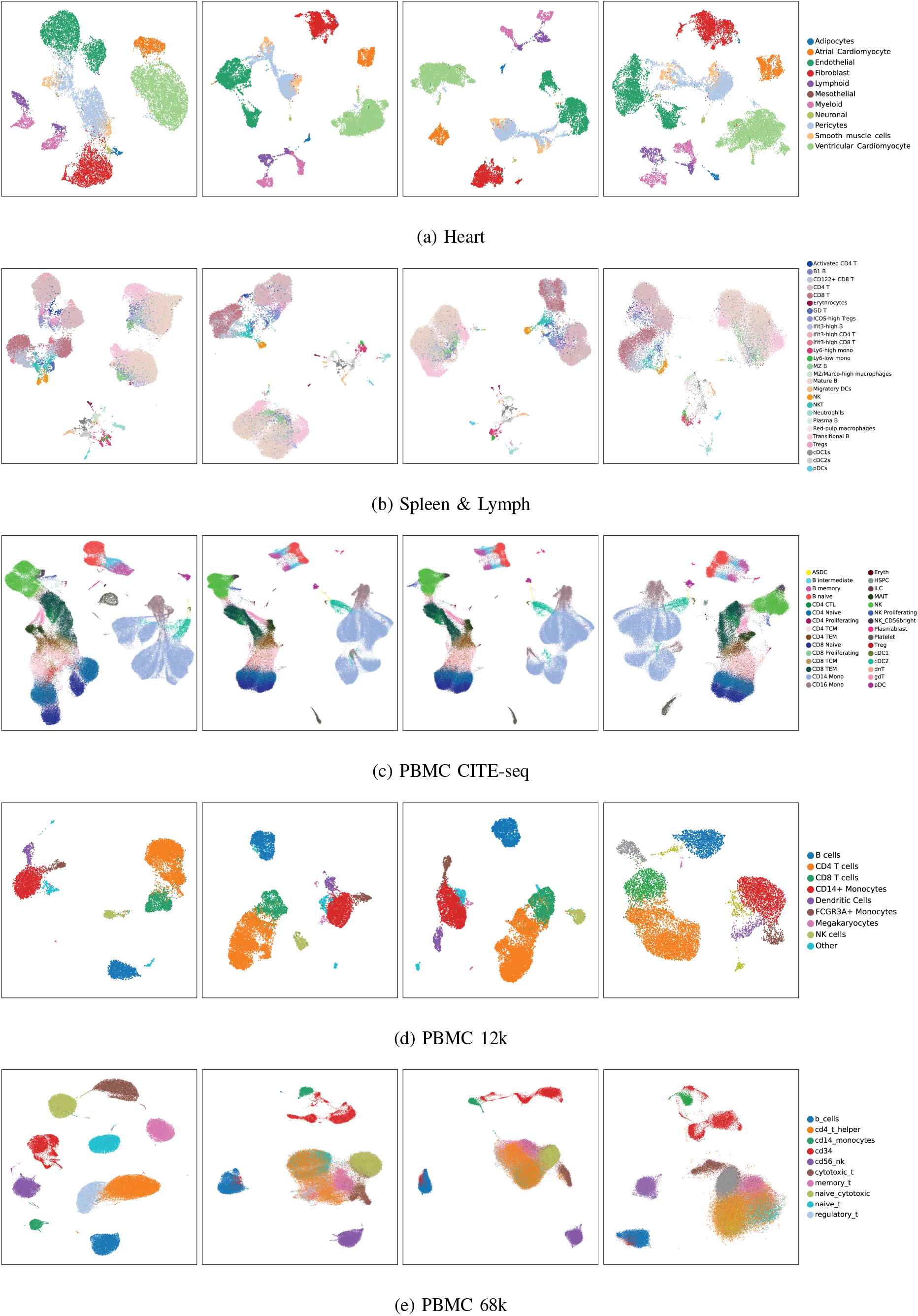
Qualitative analysis of the embedding of the proposed method, scVI, LDVAE and Scanpy from left to right, along with the ground truth labels for the corresponding datasets. Embeddings that can better separate the different ground truth labels with reduced intermixing provide better clusters with minimal overlap.

In the Heart dataset, we see that SRSA-VAE shows better cluster separation, particularly in the case of Endothelial cells. While scVI and LDVAE output more compact clusters, they overlap with the cluster of Pericyte cells. Furthermore, in both scVI and LDVAE, the Endothelial cells are also present in the cluster of ventricular cardiomyocyte cells. While scVI and LDVAE perform quite similarly, scVI performs relatively better cell type separation, which is visible when looking at the Fibroblast cells. However, SRSA-VAE outperforms the other methods since it shows minimal intermixing of different cell types in a single cluster.

The Spleen & Lymph dataset is a CITE-seq dataset, which is multi-modal in nature. This is a challenging dataset not only because of its nature but also because of the large number of cell types. The proposed method can better separate CD4T and CD8T cells. Also, B1 B cells are well separated in the proposed method.

In the case of PBMC CITE-seq datasets containing both RNA and protein expression, we tested the Spleen dataset and the PBMC CITE-seq dataset. In these datasets, SRSA-VAE showed cleaner cluster separation across multiple cell types. This is particularly visible when looking at the CD4 Naive and CD8 Naive cell clusters in the CITE-seq dataset. Furthermore, it achieves higher ARI scores in downstream clustering tasks.

For the PBMC12k dataset, we again can see better separation for the CD4T and the CD8T cells in Fig. 4d. Furthermore, other cell types are better separated from CD14+ Monocytes, which are also closely associated with FCGR3A+ Monocytes and Dendritic cells.

SRSA-VAE shows the largest margin over the competing methods on the PBMC 68k dataset, as seen in Fig. 4e. It achieves clear separation of CD4-T helper cells, Naive T cells, Memory T cells, and Naive Cytotoxic cells. In the case of scVI, LDVAE, and Scanpy, these cell types are grouped into a single cluster. All of these are subtypes of lymphoid cells and therefore share highly similar expression profiles, which collapses them under conventional encoders. The self-attention mechanism, by contextualizing each cell relative to the others in the batch, is able to resolve this substructure and yields much better cluster separation in the visualization and in downstream clustering metrics. In this case as well, scVI showed better cluster separation than its linear counterpart. We attribute this gain to the self-attention mechanism’s ability to capture subtle co-expression signatures that distinguish closely related T-cell subtypes, which conventional encoder architectures tend to collapse into a single representation. We note that the PBMC 68k ground-truth labels were originally derived via reference-based annotation against 11 purified subpopulations [14]; the exceptionally large ARI and NMI values reported for SRSA-VAE on this dataset (Tables II–III) should therefore be read as strong agreement with that reference partition rather than as a general statement of absolute performance.

**TABLE 2.**
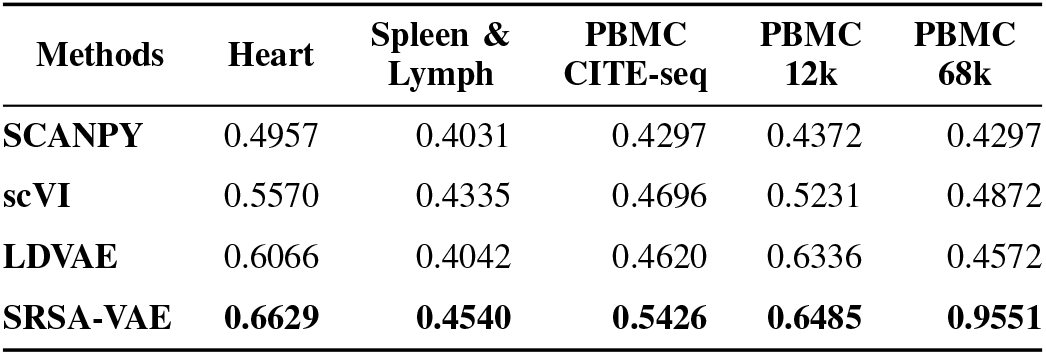
Comparison of clustering performance using Adjusted Rand Index scores for different methods.

| Methods | Heart | Spleen & Lymph | PBMC CITE-seq | PBMC 12k | PBMC 68k |
| --- | --- | --- | --- | --- | --- |
| SCANPY | 0.4957 | 0.4031 | 0.4297 | 0.4372 | 0.4297 |
| scVI | 0.5570 | 0.4335 | 0.4696 | 0.5231 | 0.4872 |
| LDVAE | 0.6066 | 0.4042 | 0.4620 | 0.6336 | 0.4572 |
| **SRSA-VAE** | **0.6629** | **0.4540** | **0.5426** | **0.6485** | **0.9551** |

**TABLE 3.** Comparison of clustering performance using Normalized Mutual Information scores for different methods.

| Methods | Heart | Spleen & Lymph | PBMC CITE-seq | PBMC 12k | PBMC 68k |
| --- | --- | --- | --- | --- | --- |
| SCANPY | 0.7298 | 0.6058 | 0.7083 | 0.7158 | 0.6045 |
| scVI | 0.7523 | 0.6273 | 0.7416 | 0.7407 | 0.6732 |
| LDVAE | 0.7764 | 0.6012 | 0.7479 | 0.7792 | 0.6476 |
| **SRSA-VAE** | **0.7875** | **0.6298** | 0.7385 | **0.7959** | **0.9592** |

#### Quantitative analysis

To quantitatively measure the clustering performance of the methods, we used Leiden clustering on the latent embedding and then measured the ARI, NMI and V-measure scores for each method. We also measured the Silhouette score and Calinski-Harabasz score.

SRSA-VAE performs well on the benchmark scRNA-seq datasets as seen in Table II and Table III. The ARI scores on the Heart, PBMC 12k, and PBMC 68k are considerably higher than the state-of-the-art methods. The gains on the CITE-seq datasets, that is on the Spleen & Lymph dataset and the PBMC CITE-seq dataset, are more modest but still competitive. NMI scores follow a broadly similar trend, with SRSA-VAE achieving the highest NMI on Heart, Spleen & Lymph, PBMC 12k, and PBMC 68k; on the PBMC CITE-seq dataset, SRSA-VAE NMI is comparable to the best competing method.

In Table IV, we see that SRSA-VAE scores are better in terms of homogeneity scores. However, it marginally falls behind in a few cases when it comes to completeness scores. However, a lower completeness score means that the clustering algorithm partitioned a single cluster into multiple clusters. This can be beneficial for cell sub-population discovery. This is further corroborated by Figure 3. In PBMC CITE-seq and the PBMC 12k, the proposed method generated more clusters.

**TABLE 4.** Comparison of homogeneity, completeness and v-measure scores for the different methods.

|  | Methods | Heart | Spleen & Lymph | PBMC CITE-seq | PBMC 12k | PBMC 68k |
| --- | --- | --- | --- | --- | --- | --- |
| Homogeneity | SCANPY | 0.911 | 0.492 | 0.673 | 0.892 | 0.637 |
|  | scVI | 0.892 | 0.598 | 0.658 | 0.838 | 0.670 |
|  | LDVAE | 0.907 | 0.582 | 0.665 | 0.847 | 0.673 |
|  | SRSA-VAE | 0.887 | 0.610 | 0.794 | 0.892 | 0.978 |
| Completeness | SCANPY | 0.587 | 0.446 | 0.714 | 0.598 | 0.576 |
|  | scVI | 0.674 | 0.618 | 0.736 | 0.663 | 0.658 |
|  | LDVAE | 0.658 | 0.602 | 0.800 | 0.722 | 0.650 |
|  | SRSA-VAE | 0.707 | 0.641 | 0.690 | 0.719 | 0.941 |
| V-measure | SCANPY | 0.714 | 0.468 | 0.693 | 0.716 | 0.605 |
|  | scVI | 0.768 | 0.608 | 0.695 | 0.741 | 0.664 |
|  | LDVAE | 0.762 | 0.592 | 0.726 | 0.779 | 0.662 |
|  | SRSA-VAE | 0.787 | 0.625 | 0.738 | 0.796 | 0.959 |

We also explore the clustering performance using intrinsic clustering metrics. The silhouette score provides a quantitative measure of cluster cohesion and separation. Across all the datasets, we see marginal gains by using the proposed method as seen in Table V. Moreover, we also use the Calinski-Harabasz score as another metric for assessing cluster quality. Across all the datasets, the proposed method outperforms other methods, exhibiting a high between-cluster variance and low within-cluster variance.

**TABLE 5.** Silhouette score and Calinski-Harabasz score for the different methods on five datasets.

|  | Methods | Heart | Spleen & Lymph | PBMC CITE-seq | PBMC 12k | PBMC 68k |
| --- | --- | --- | --- | --- | --- | --- |
| Silhouette Score | SCANPY | 0.2029 | 0.027 | 0.082 | 0.200 | 0.090 |
|  | scVI | 0.2269 | 0.083 | 0.141 | 0.244 | 0.095 |
|  | LDVAE | 0.2728 | 0.082 | 0.112 | 0.279 | 0.134 |
|  | SRSA-VAE | **0.3263** | **0.087** | **0.294** | **0.377** | **0.619** |
| Calinski-Harabasz Score | SCANPY | 2014.616 | 1768.209 | 9837.172 | 1854.285 | 8246.119 |
|  | scVI | 1666.628 | 1034.584 | 9005.577 | 1445.262 | 8433.480 |
|  | LDVAE | 2595.812 | 1249.447 | 8407.534 | 1526.875 | 11565.113 |
|  | SRSA-VAE | **12976.324** | **1796.209** | **51266.863** | **3879.513** | **167021.680** |

### B. Analysis of the model stability and parameter setting

For dimensionality reduction using autoencoders, parameter tuning is an essential part [20]. In the proposed method, the parameters that had the maximum effect on the performance of the downstream clustering task are the number of epochs and the dimension of latent variables. Both the parameters were fixed using a grid-search mechanism.

To determine the number of epochs, we used an early stopping mechanism where the model stops training after the validation loss does not decrease in 10 epochs. We found that training the proposed model for less than 25 epochs usually provided the best performance in clustering. Figure 6 shows the training loss and the validation loss of the models across the five datasets along with the ARI of the clustering. It is observed that most models converge within 100 epochs whereas the ARI value peaks during the earlier epochs.

The number of latent variables also has a notable impact on the clustering performance. We tested the models to generate the latent embedding with sizes varying from 5 to 12. This is shown in Figure 5. The heart dataset provides the best performance when the size of the latent variable is set to 10. The same is followed by the PBMC 68k dataset. The Spleen & lymph dataset performs best with a 9-dimensional latent embedding, whereas PBMC CITE-seq does best with 8 and PBMC 12k does best with 7. Increasing the number of latent variables beyond 12 further reduced the clustering accuracy.

**Fig. 5.**
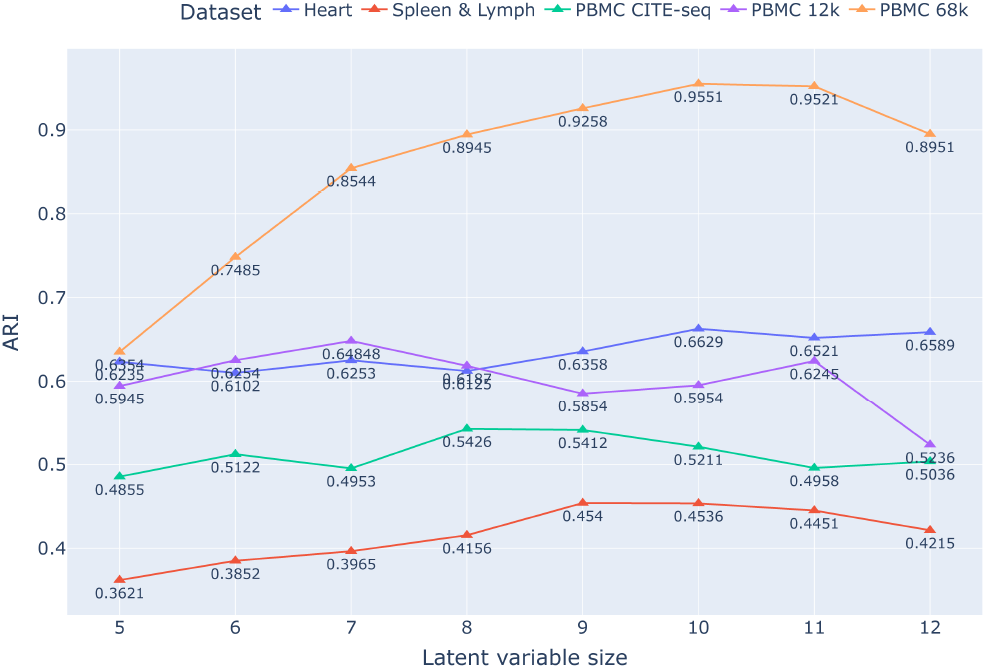
Variation of clustering accuracy with the change of the size of the latent variable.

**Fig. 6.**
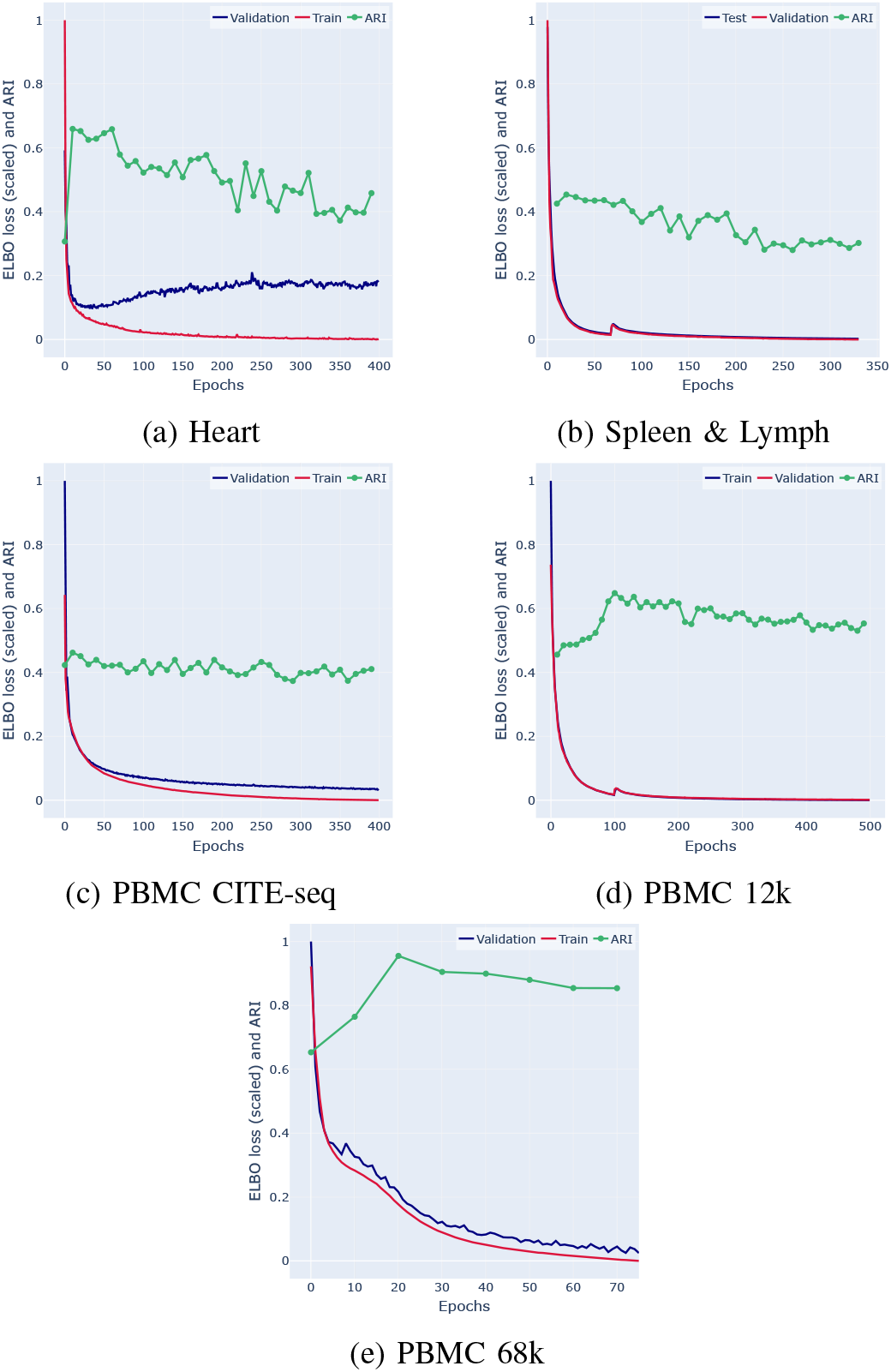
Training and validation losses versus the Adjusted Rand Index for the proposed model on five datasets.

### C. Effectiveness of Residual Connection and Attention mechanisms

To understand the effectiveness of the novel encoder mechanism, we performed a thorough ablation study to verify the effectiveness of the self-attention mechanism and the scalable residual connection. The observations of the ablation study are presented in Table VI.

**TABLE 6.** Ablation study showing the effect of the self-attention mechanism and the scalable residual connection on the clustering performance.

| Method | Heart | Spleen & Lymph | PBMC CITE-seq | PBMC 12k | PBMC 68k |
| --- | --- | --- | --- | --- | --- |
| **Proposed method** | 0.6629 | 0.454 | 0.5426 | 0.6484 | 0.9551 |
| **Without scalable-residual connection** | 0.5923 | 0.3823 | 0.4928 | 0.5454 | 0.7845 |
| **Without self-attention** | 0.6256 | 0.4156 | 0.5158 | 0.5394 | 0.8207 |

The variant without the scalable residual connection simply has the self-attention mechanism between the initial and the final linear layers. However, this seems to be detrimental to performance since self-attention layers can cause overfitting [21]. Hence, we used a residual connection to make the training easier and to preserve the input data. This creates latent embedding that is better for the downstream clustering tasks.

The variant without the self-attention mechanism simply replaces the self-attention mechanism with a single linear layer. This makes the encoder a simple three-layer neural network with a residual connection from the first layer to the third layer. The *α* and *β* parameters are preserved in this variant and they are kept trainable. The input to the first layer is concatenated with the output of the second layer and is then fed to the third layer. Without the self-attention mechanism and only with the scalable residual mechanism provides marginally better performance. However, combining both mechanisms provides the best performance.

### D. Comparison with the model for Multimodal Data

To evaluate performance on multimodal data, we compare SRSA-VAE with TotalVI [5], a model built on the scVI framework for integrating RNA and protein expression in CITE-seq datasets. TotalVI uses separate encoders for gene and protein modalities, combining them into a unified latent space for downstream tasks such as clustering and visualization.

We assess the encoder components of both models on two CITE-seq datasets: PBMC and Spleen and Lymph. In both cases, latent embeddings are extracted and clustered using the Leiden algorithm following Scanpy’s standard workflow. Our results in Table VII show that SRSA-VAE achieves consistently higher ARI scores than TotalVI, indicating improved clustering accuracy. This suggests that a unified self-attention-based encoder can more effectively integrate multimodal information than modality-specific encoders. Additionally, the inclusion of a scalable residual connection in SRSA-VAE further boosts performance, particularly on large datasets.

**TABLE 7.**
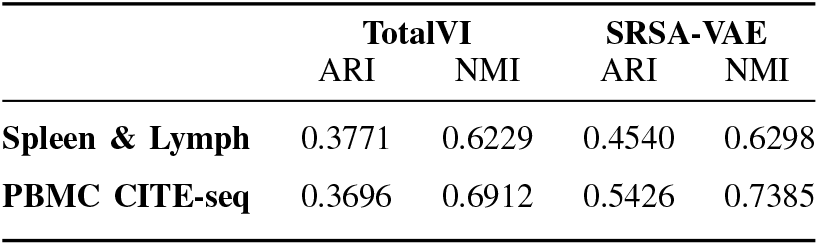
Comparison with TotalVI is also performed to show the advantage of using self-attention to create a unified representation of protein and RNA representations instead of using separate encoding mechanisms.

|  | TotalVI |  | SRSA-VAE |  |
| --- | --- | --- | --- | --- |
|  | ARI | NMI | ARI | NMI |
| **Spleen & Lymph** | 0.3771 | 0.6229 | 0.4540 | 0.6298 |
| **PBMC CITE-seq** | 0.3696 | 0.6912 | 0.5426 | 0.7385 |

### E. Runtime analysis

All models were trained on Google Colab Pro+ instances with 16GB NVIDIA V100 GPUs and 64GB RAM. For fair comparison, SRSA-VAE was trained with a fixed 100-epoch limit and early stopping based on validation performance. Unlike scVI, which uses a dataset-dependent heuristic to set epochs, SRSA-VAE dynamically halts training to minimize runtime. Although its per-epoch cost is higher, the total training time remains significantly lower. As shown in Figure 7, SRSA-VAE achieves competitive ARI scores while requiring substantially less training time, demonstrating its efficiency and suitability for clustering large-scale single-cell datasets.

**Fig. 7.**
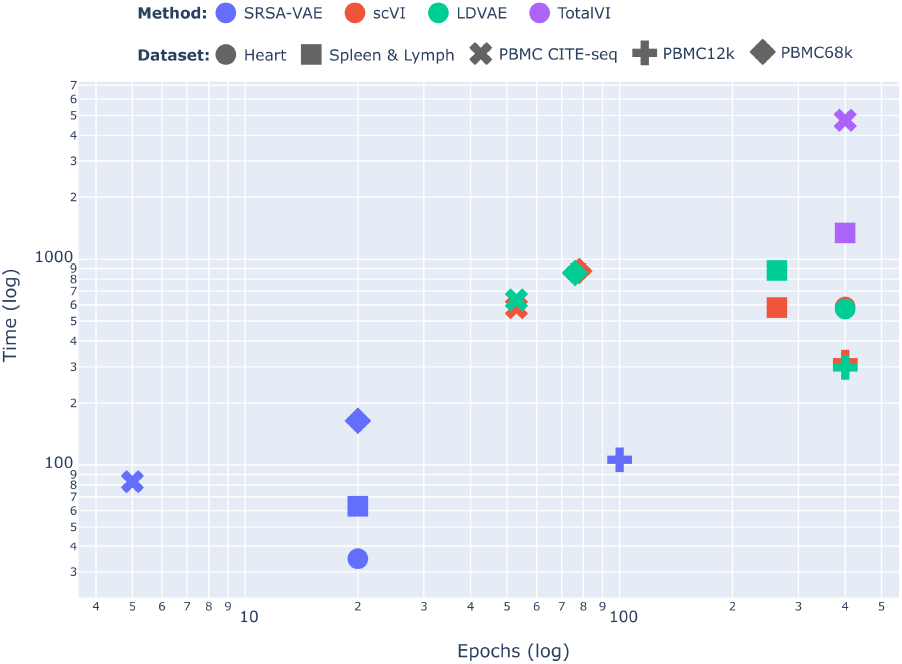
Time required by the various autoencoder-based methods to generate features to achieve optimal clustering. While SRSA-VAE requires marginally more time per epoch, the drastically reduced number of epochs shortens the required training time significantly.

## V. Conclusion

We proposed SRSA-VAE, a variational autoencoder with a scalable residual self-attention encoder for feature learning in single-cell RNA-seq and CITE-seq data. Unlike existing autoencoder-based methods that optimize purely empirical loss functions, SRSA-VAE formalizes an attention-weighted variational inference mechanism that explicitly encodes feature dependencies into the latent space, offering a theoretical advancement over standard VAEs and their multimodal extensions. These design choices yield compact, informative embeddings that improve clustering performance. Ablation studies confirm that both the attention mechanism and residual connection significantly impact performance. The model converges faster and achieves high Adjusted Rand Index and silhouette scores across diverse datasets, including multimodal data. Although we compare against established models that offer more comprehensive functionality beyond clustering, these models often require extensive pre-training or are designed for batch correction and trajectory inference. In future work, we plan to evaluate SRSA-VAE in such settings and extend its application to unseen tissue types.

## Data Availability

All datasets used in this study are publicly available. The adult human heart scRNA-seq data are available from Litviňuková *et al*. [11]. The murine Spleen and Lymph Node CITE-seq data are distributed with totalVI [5]. The PBMC CITE-seq multimodal reference atlas is available from Hao *et al*. [12]. The PBMC 12k (4k + 8k) scRNA-seq data are available from the 10x Genomics public datasets repository [13]. The PBMC 68k scRNA-seq data are available from Zheng *et al*. [14]. No new data were generated in this study.

## Code Availability

The reference implementation of SRSA-VAE, together with the training and evaluation scripts used to produce the results in this paper, is openly available under an open-source license at https://github.com/rangan2510/srsa-vae.

## Author Contributions

R.D. and A.D. contributed equally to this work. U.M. and S.B. supervised the project. All authors reviewed and approved the final manuscript.

## Competing Interests

The authors declare no competing interests.

## Acknowledgment

RD, UM, and SB acknowledge the support from CEFIPRA, Project No. 6702-1. AD and SB acknowledge the Science and Engineering Research Board (SERB), Government of India for providing the JC Bose Fellowship and research grant (JBR/2021/000036).

## Notes

https://github.com/rangan2510/srsa-vae

### Competing Interest Statement

The authors have declared no competing interest.

